# Structural variation discovery in wheat using PacBio high-fidelity sequencing

**DOI:** 10.1101/2023.12.08.570887

**Authors:** Zhiliang Zhang, Jijin Zhang, Lipeng Kang, Xuebing Qiu, Song Xu, Jun Xu, Yafei Guo, Zelin Niu, Beirui Niu, Aoyue Bi, Xuebo Zhao, Daxing Xu, Jing Wang, Changbin Yin, Fei Lu

## Abstract

**Background:** Structural variations (SVs) pervade plant genomes and contribute substantially to the phenotypic diversity. However, most SVs were ineffectively assayed because of their complex nature and the limitations of early genomic technologies. The recent advance in third-generation sequencing, particularly the PacBio high-fidelity (HiFi) sequencing technology, produces highly accurate long-reads and offers an unprecedented opportunity to characterize SVs’ structure and functionality. As HiFi sequencing is relatively new to population genomics, it is imperative to evaluate and optimize HiFi sequencing based SV detection before applying the technology at scale.

**Results:** We sequenced wheat genomes using HiFi reads, followed by a comprehensive evaluation of mainstream long-read aligners and SV callers in SV detection. The results showed that the accuracy of deletion discovery is markedly influenced by callers, which account for 87.73% of the variance, while both aligners (38.25%) and callers (49.32%) contributed substantially to the accuracy variance for insertions. Among the aligners, Winnowmap2 and NGMLR excelled in detecting deletions and insertions, respectively. For SV callers, SVIM achieved the best performance. We demonstrated that combining the aligners and callers mentioned above is optimal for SV detection. Furthermore, we evaluated the effect of sequencing depth on the accuracy of SV detection, showing that low-coverage HiFi sequencing is sufficiently robust for high-quality SV discovery.

**Conclusions:** This study thoroughly evaluated SV discovery approaches using HiFi reads, establishing optimal workflows to investigate structural variations in the wheat genome. The notable accuracy of SV discovery from low-coverage HiFi sequencing indicates that skim HiFi sequencing is effective and preferable to characterize SVs at the population level. This study will help advance SV discovery and decipher the biological functions of SVs in wheat and many other plants.

## Background

Structural variations (SVs) and single nucleotide polymorphisms (SNPs) are at the opposite ends of the genetic variation spectrum. Contrary to the simplicity of SNPs, SVs exhibit a much higher level of complexity—insertion, deletion, duplication, inversion, and translocation, varying in size from ∼50 bp [1] to hundreds of megabases (Mb) [2], constitute a highly diverse set of genetic variations in the genome [3, 4]. In contrast to the human genome, where SVs affect a mere 1.5% of the genome when comparing two individuals [3, 4], the plant genome is in flux—due to frequent genome duplications and the proliferation of transposable elements (TEs) [5], SVs can comprise up to 50% of the genome when comparing two individuals in one species [6–8]. A growing body of genomic analyses has shown the pivotal role of SVs in shaping plant phenotypic traits, with a broad scope from gene expression to environmental adaptation [9–14]. Meanwhile, quantitative trait locus (QTL) mapping has identified SVs as causative genetic variations controlling important agronomic traits of crops, such as flowering time in maize (e.g., *Vgt1* [15]*, ZmCCT* [16, 17]), grain yield in rice (e.g., *GW5* [18] and *GL7* [19]), solid-stemmed architecture in wheat (e.g., *TdDof* [20]), and aroma volatiles in tomato (e.g., *NSGT1* [21] and *NSGT2* [9]), etc. Despite the recognized prevalence and importance of SVs in plant genomes, accurate genotyping of SVs remains a substantial challenge in genome decoding efforts [22–25].

Aside from the intrinsic structural complexity of SVs, technological limitations are impeding effective SV detection and genotyping [4, 26]. Although the cost of next-generation sequencing (NGS) keeps declining, the effectiveness of this technology is constrained by the short read length, which is inadequate to detect large SVs. The initial form of long-read sequencing, such as PacBio continuous long reads (CLR) and Oxford Nanopore, provides a solution for identifying large SVs, albeit with a high sequencing error rate (∼8-13%) [27] that may lead to an inaccurate discovery of small SVs (100-200bp) [28] and the imprecise detection of SV breakpoint [29, 30]. Encouragingly, the new PacBio Circular Consensus Sequencing (CCS, a.k.a. HiFi sequencing) strikes the perfect balance between low error rate (< 0.2%) and long read length (> 10 kb), resulting in much-improved performance on SV detection [31]. Recently, a growing number of long-read algorithms, including both aligners (e.g., pbmm2, NGMLR [31], Winnowmap2 [32], and Minimap2 [33]) and SV callers (e.g., pbsv, cuteSV [34], SVIM [35], SVDSS [36], and Sniffles2 [37]), have been developed for SV detection [38, 39]. With the great stride in sequencing technology and toolset development, it is important to benchmark bioinformatic tools to optimize the use of HiFi reads while performing SV detection in plant genomes.

Bread wheat is one of the most important crops worldwide. Meanwhile, it is a hexaploid species with a genome of ∼85% TE content [40], well exemplifying the intricate nature of plant genomes. This study sequenced wheat accessions and performed a comprehensive evaluation of HiFi sequencing for SV detection in plant genomes. Through the assessment of mainstream long-read aligners and SV callers, our analysis provides the optimal workflow of SV genotyping with HiFi reads, which is anticipated to help dissect SVs’ functionality in wheat and many other plants.

## Results

### Overview of developing the SV truth set

A truth set of SVs is essential to benchmark SV discovery approaches. Several projects, including Genome in a Bottle (GIAB) [41] and Human Genome Structural Variation Consortium (HGSV) [42], have provided the SV reference data sets for human genetic studies. However, such SV truth sets are lacking for wheat. To address this, we embarked on constructing a robust wheat SV truth set from the ground up, a process consisting of three steps (Fig. 1). First, we generated comprehensive SV call sets, which were expected to include as many candidate SVs as possible. The goal was achieved by calling SVs through applying pairwise combination of long-read aligners and SV callers on HiFi reads. Second, we constructed the SV truth set by verifying all candidate SVs using high-depth NGS. Despite the limitation in identifying the full structure of large SVs, NGS provides precise alignment signatures at SV breakpoints [28], thereby offering strong validation for SVs initially detected by HiFi reads. Third, we applied costly or low-throughput, but independent and proven approaches, such as de novo assembly, PCR amplification, and Sanger sequencing, to evaluate the integrity of the SV truth set.

**Fig. 1.**
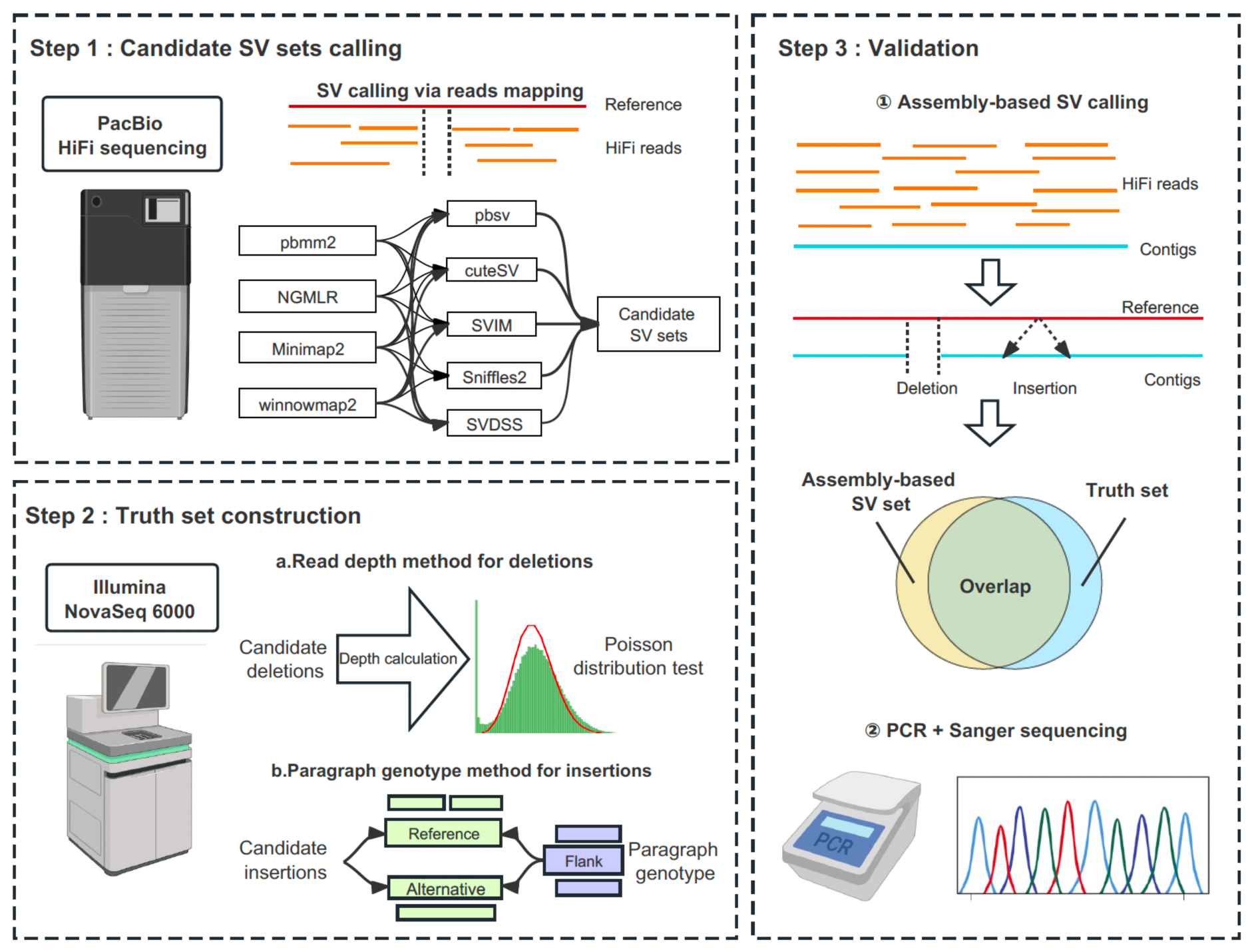
Overview of developing the truth set. **Step 1: Candidate SV sets calling.** Candidate SV sets were generated from the combination of long-read aligners and SV callers based on PacBio HiFi reads. **Step 2: Truth set construction.** SV truth set was formed by verifying all candidate SVs using deep-coverage next-generation sequencing. **Step 3: Validation of the truth set.** The truth set was validated by assembly-based SV calling, PCR amplification, and Sanger sequencing.

### The sequence-resolved SV truth set

Bread wheat (*Triticum aestivum* ssp. *aestivum*, 2*n* = 6*x* = 42, AABBDD) originated from the natural polyploidization event between the free-threshing tetraploid wheat (*Triticum turgidum*, 2*n* = 4*x* = 28, AABB) and the diploid goatgrass (*Aegilops tauschii* ssp. strangulata, 2*n* = 2*x* = 14, DD) around ∼10,000 years ago [43]. Frequent gene flow resulted in both shared and distinct genetic diversities among bread wheat and its polyploid progenitors [44]. To ensure that our evaluation is equally useful for the entire gene pool of bread wheat, we sequenced three plant accessions from bread wheat, wild emmer wheat (*Triticum turgidum* ssp. *dicoccoides*), and strangulata (*Aegilops tauschii* ssp. *strangulata*). Each sample was sequenced with one SMRT cell using the PacBio CCS mode, achieving individual coverage of 1.7×, 2.7×, and 6.6×. These highly accurate HiFi reads (accuracy >99.9%) reached an average length of 13.0 kb, 17.2 kb, and 12.9 kb, respectively (Supplementary Table 1 and Supplementary Fig. 1).

We performed the candidate SV discovery by applying a full combination of leading long-read aligners (pbmm2, NGMLR, Winnowmap2, and Minimap2) and SV callers (pbsv, cuteSV, SVIM, SVDSS, and Sniffles2), yielding 20 distinct sequence-resolved SV call sets from the aligner-caller-combinations (ACCs). The SV call sets from different ACCs varied in number and size (Fig. 2). We observed that the SV count derived from SVDSS was consistently lower than other callers, which was 38.45%-43.21% for deletion and 24.00%-29.77% for insertion when compared to results from pbsv, cuteSV, SVIM, and Sniffles2. Meanwhile, these call sets presented an approximately 8-fold change in average SV length across different ACCs, with deletions ranging from 273 bp to 1,523 bp and insertions from 182 bp to 1,853 bp, respectively (Supplementary Table 2).

**Fig. 2.**
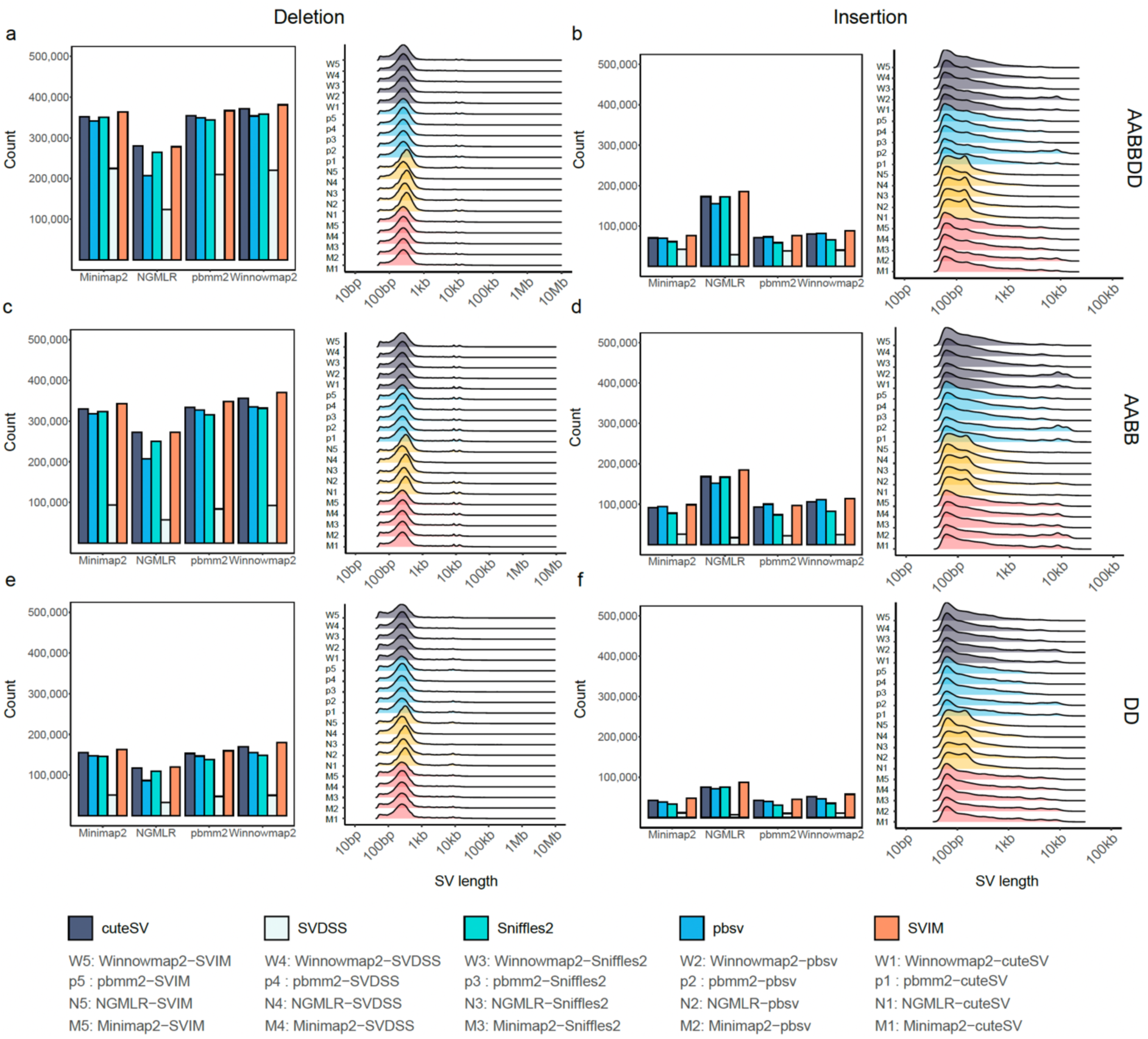
The sequence-resolved candidate SV sets of 20 aligner-caller-combinations (ACCs). The count and length distribution of deletion (**a, c, e**) and insertion (**b, d, f**) from 20 sequence-resolved candidate SV sets in three wheat accessions of AABBDD, AABB, and DD genomes.

We used high-depth NGS to discern true SVs from the 20 SV call sets. The three wheat accessions mentioned above were whole-genome sequenced with read depths of 17×, 14×, and 25×, respectively. For deletion, the abnormal low read depth was used to distinguish true deletion from candidates (see Methods). For insertion, we used paragraph [45], which leverages discordant alignment features (e.g., read depth, read pair, and split read), to identify true insertions (see Methods). This rigorous verification process yielded a refined SV truth set comprising 1,030,303 deletions and 1,457,547 insertions, with average lengths of 1,375 bp and 571 bp, respectively (Supplementary Table 3). The multi-modal size distribution of the SV truth set indicates the aggregation of candidate SVs from multiple ACCs (Fig 2 and Fig. 3a,d). We found that these true SVs were distributed along entire chromosomes, covering both distal chromosomes and pericentromeric regions, showing that the truth set is qualified to benchmark SV call sets at the whole-genome level (Fig. 3b,e).

**Fig. 3.**
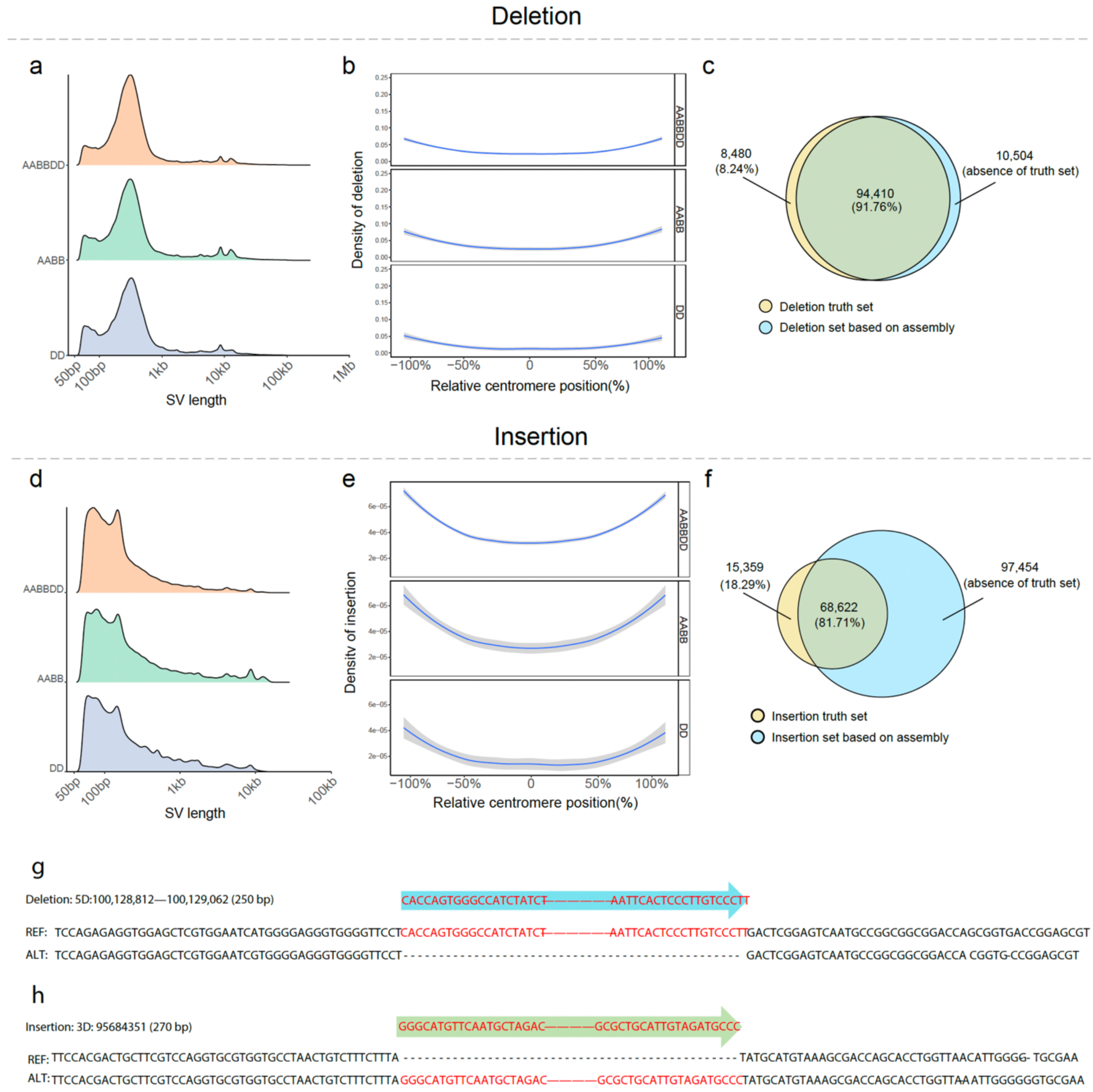
Validation of the SV truth set. The count **(a, d**) and genome distribution **(b, e**) of deletion and insertion truth set. A total of 91.76% of deletions and 81.71% of insertions in the SV truth set were validated by assembly-based SV call set **(c, f**). Sanger sequencing results showcased two examples of 250 bp deletion and 270 bp insertion with precise SV breakpoint (**g, h**).

To evaluate the integrity of the SV truth set, we implemented two independent approaches for the assessment. First, we performed an assembly-based validation. As the diploid goatgrass (DD genome) was sequenced with HiFi reads at the coverage of ∼6.6×, we de novo assembled the long-read data into contigs using HiCanu [46]. The assembly contained 36,983 contigs with contig N50 of 164 kb. A total of 316,038 SVs, including 162,784 deletions and 153,254 insertions, were discovered by mapping the contigs to the wheat reference genome (see Methods). We found that 91.76% of deletions and 81.71% of insertions in the SV truth set were confirmed by the assembly-based SV call set (Fig. 3c,f). Second, we employed PCR amplification and Sanger sequencing to verify SVs with the length ranging from 200 bp to 600 bp (Supplementary Fig. 2). Ten deletions and ten insertions, chosen at random, were all successfully amplified by PCR (Supplementary Fig. 2). Followed Sanger sequencing results showed marked accuracy of SV breakpoints and corroborated the SV sequences in the truth set (Fig. 3g,h, Supplementary Fig. 3-5, and Supplementary Table 5). Both lines of evidence demonstrate the high accuracy of the sequence-resolved SV truth set.

### Performance comparison of aligner-caller-combinations

We then benchmark the 20 ACCs by comparing the corresponding SV call set against the ground truth. The results showed that the ACCs had different patterns of performance while detecting deletions and insertions. These ACCs exhibited higher precisions (one-tailed *t-*test, *P*-value = 1.53×10^-10^) and recalls (one-tailed *t-*test, *P*-value = 6.39×10^-5^) in discovering deletions compared to insertions. Regarding variability, these ACCs showed similar precisions (SD = 0.01) but varied recalls (SD = 0.20) while detecting deletions. In contrast, they showed considerable variation in both precisions (SD = 0.07) and recalls (SD = 0.21) when it came to detecting insertions. To evaluate the effects of aligners and callers on the accuracy of SV calling, we conducted an analysis of variance (ANOVA) of precision, recall, and F-score using the benchmarking results. We found that callers predominantly accounted for the variance in accuracy for deletions, while both aligners and callers contributed substantially to the accuracy variance for insertions. Taking the F-score for example, the callers explained 87.73% (*F*-test, *P*-value = 3.57×10^-11^) of variance and aligners explained only 11.10% (*F*-test, *P*-value = 2.11×10^-6^) while detecting deletions. Conversely, the callers explained 49.32% (*F*-test, *P*-value = 3.86×10^-4^) of variance and aligners explained 38.25% (*F*-test, *P*-value = 5.67×10^-4^) while detecting insertions. These results suggest that the accuracy of detecting deletions is largely attributed to the sensitivity of callers, and the accuracy of detecting insertions depends on both the sensitivity and specificity of aligners and callers.

Compared to the individual methods, the ensemble approach, which combines SV call sets from multiple methodologies through intersection or union, has demonstrated enhanced precision or recall in SV detection using NGS [47, 48]. To explore the potential benefits of the ensemble approach with HiFi reads, we integrated multiple SV call sets of ACCs and benchmarked their collective performance. By randomly sampling 20 ACCs and combing their SV call sets, we created subgroups of ACCs with varying size from 2 to 20, and generated 1,600 ensemble SV sets through intersecting and uniting SV call sets of ACCs (see Methods, Supplementary Table 7). The benchmarking results showed that the intersection of deletion call sets increased precision at the cost of recall, whereas the union of deletion call sets exhibited the reverse tendency. The integration of more ACCs into the ensemble intensified these trends (Fig. 4b, Supplementary Fig. 7-9b). Similarly, for insertions, intersection led to greater precision but lower recall. Interestingly, unlike deletions, both precision and recall were improved when insertion call sets were united (Fig. 4e, Supplementary Fig. 7-9e).

**Fig. 4.**
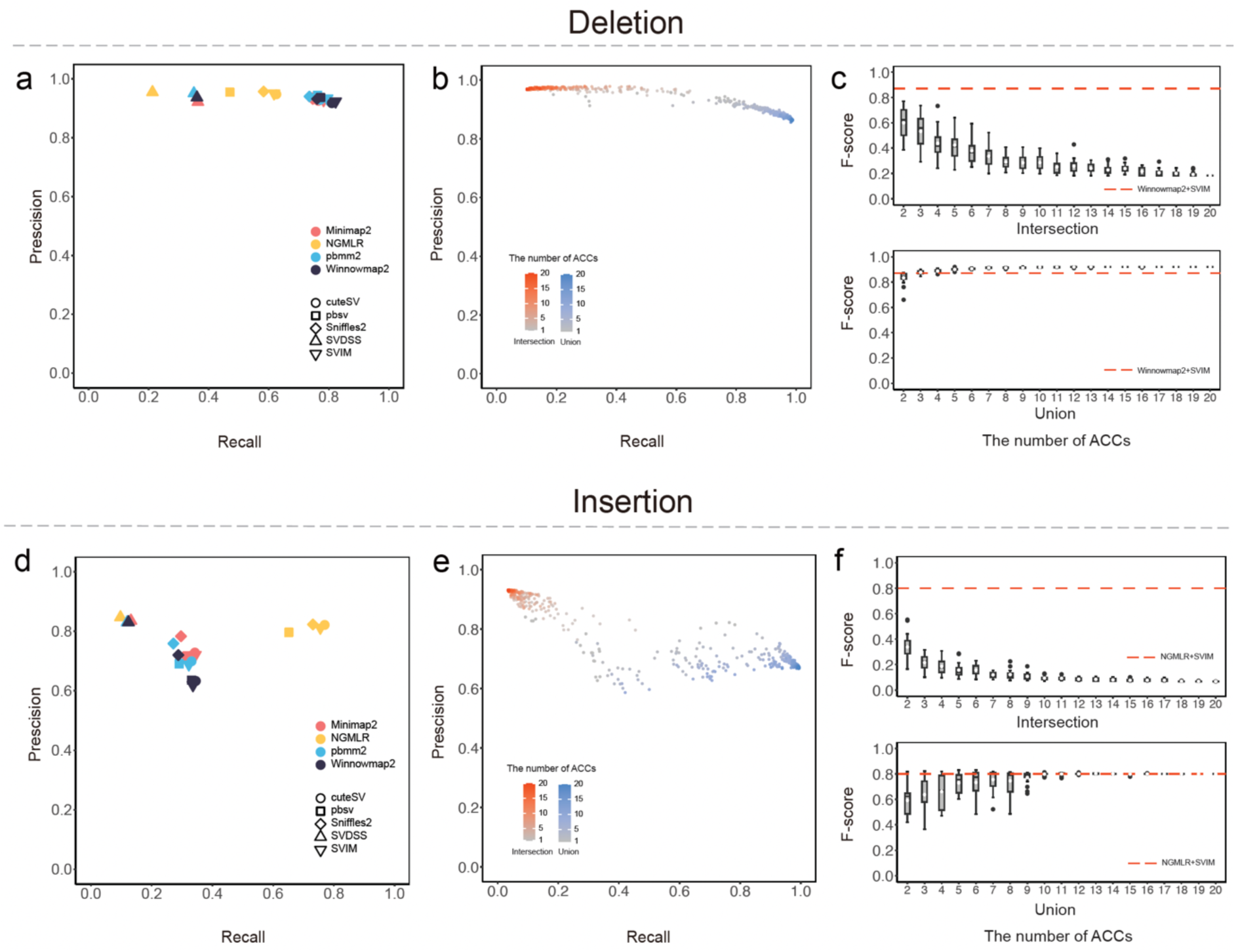
Evaluation of individual ACCs and the ensemble approach. The precision-recall graph of 20 ACCs against the SV truth set (**a, d**). The performance of 1,600 ensemble SV sets through intersecting and uniting SV call sets of ACCs (**b, e**). The F-score comparison between 1,600 ensemble SV sets and the individual call set with the highest F-score in 20 ACCs (**c, f**). The individual method with highest F-score of 20 ACCs was represented by red dashed line.

Upon evaluating the F-score, we found that of the individual method showed superb performance when compared with the ensemble approach. For deletion, the F-score of the Winnowmap2-SVIM call set outperformed all the intersection sets and was close to the F-score plateau of union sets (Fig. 4c, Supplementary Fig. 7-9c). For insertion, the NGMLR-SVIM call set surpassed all intersection sets and its F-score was nearly identical to the plateau value of union sets (Fig. 4f, Supplementary Fig. 7-9f). Although a number of ensemble SV call sets exhibited higher precision or recall, we consider that the optimal individual method is efficient, given that ensemble approach presents a clear precision-recall trade-off (Fig. 4b,e) and it demands considerably more computational time and data storage (Supplementary Table 8).

### Low-coverage HiFi sequencing for SV discovery

As there is a recognized trade-off between sequencing coverage and SNP calling accuracy, adequate coverage was considered essential to ensure the SNP accuracy using NGS [49]. However, given that the random insertion error in previous PacBio long reads has been addressed in HiFi sequencing [39, 50], we speculated that the coverage of HiFi sequencing may not strongly affect the false positive rate of SV calling. To test the hypothesis, we downsampled the HiFi reads of strangulata, the diploid wheat accession that was sequenced at the coverage of 6.6×, to examine the performance of the 20 ACCs at different levels of sequencing depth. By benchmarking these SV call sets against the truth set, the results showed that the recall improved with the increasing sequencing depth, but with a clear diminishing return irrespective of the aligner or caller used (Fig. 5a,c and Supplementary Table 9). However, precision did not increase with higher coverage for all the 20 ACCs. On the contrary, these ACCs showed a slight decline in precision as the coverage increased (Fig. 5b,d and Supplementary Table 9), which is probably due to the inconsistency of alignment of individual reads. These findings suggest that while low-coverage HiFi sequencing might be prone to technological missing data, it does not compromise the validity of detected SVs.

**Fig. 5.**
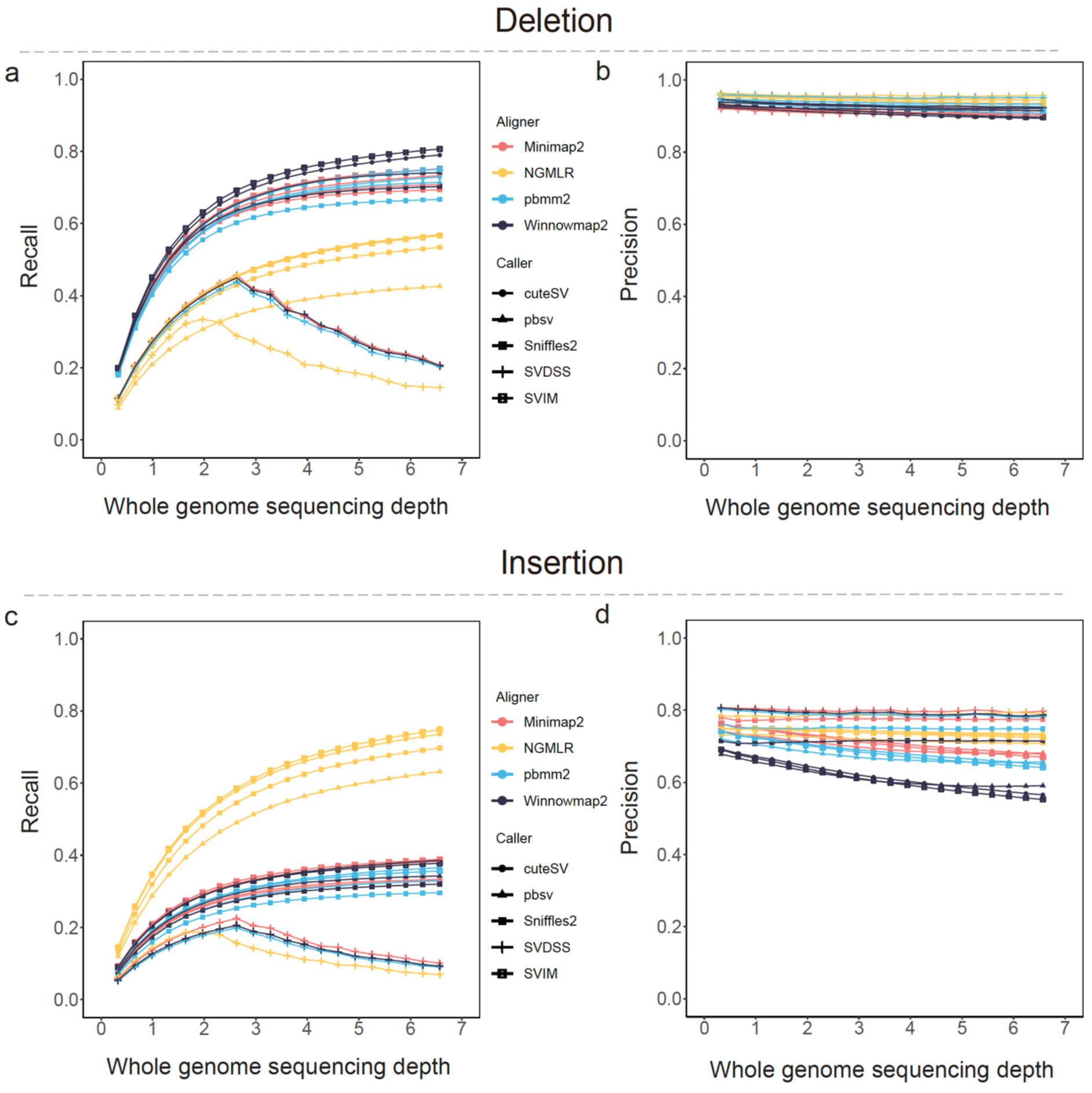
Evaluation of low-coverage HiFi sequencing for SV discovery. Precision (**a, c**) and recall (**b, d**) showed the effect of the sequencing depth from 0.33X to 6.60X.

## Discussion

Plant genomes are incredibly complex, where an immense number of SVs are regulating various phenotypic traits. While historically challenging, functional analysis of SVs has started taking off because of recent breakthroughs in long-read sequencing [9, 51–59]. As the latest long-read technology, HiFi sequencing has markedly improved read accuracy to >99.8% [50] and will likely become increasingly important in SV discovery [39, 60, 61]. Concurrently, new SV detection algorithms, including both long-read aligners and SV callers, are under rapid development [38, 39]. Given these advancements, there is a pressing necessity to assess the performance of these bioinformatic tools, especially for the intricate plant genomes. Our study timely addressed the need by sequencing wheat accessions at the whole-genome scale, establishing a sequence-resolved SV truth set, and conducting an exhaustive evaluation of the SV discovery methods that leverage HiFi sequencing.

Different from previous studies that focus solely on callers when analyzing SV discovery methods using NGS [47, 48], our study extends the evaluation to involve both aligners and callers. The analysis utilizing HiFi reads revealed distinct contributions to the variance of the F-score—callers accounted for 87.73% when identifying deletions, while aligners contributed a minor portion of 11.10%; in contrast, for insertions, callers were responsible for 49.32% of the variance, and aligners for 38.25% (Supplementary Table 6). The sharp difference in the F-score’s variance components for deletions and insertions, along with the superior F-score observed in deletion detection (Fig. 4a,d), underscores the inherent difficulties in identifying insertions within the genome. These findings also suggest that adopting a tailored approach, which employs the most efficacious methods for detecting each SV type separately, could be a good practice to enhance the accuracy of SV discovery in genomic studies.

Previous analyses have indicated that combining multiple SV call sets derived from either NGS [29] or Oxford Nanopore Technologies (ONT) data [30] can improve the accuracy. In pursuit of an effective SV discovery workflow, our study aggregated 20 individual SV call sets from ACCs and created 1,600 ensemble SV sets by intersecting and uniting these ACCs for the detection of deletions and insertions, respectively (Fig. 4). The benchmarking results showed that integrating more ACCs led to increased confidence (through intersection) and sensitivity (through union) in general (Fig. 4b,e). Notably, the individual method with the highest F-score demonstrated sufficient efficiency and outperformed most of the ensemble SV sets, challenging the assumption that ensemble methods invariably exceed the individual method. This result is consistent with an earlier evaluation of SV callers showing that “Ensemble calling is not a panacea.” [47] This finding, coupled with the significant computational resources and data storage required for the ensemble approach, highlights the utility of the individual approach. Specifically, the caller SVIM, when used after Winnowmap2 for deletion detection or NGMLR for insertion detection, stands out as an effective and resource-efficient choice. Hence, we recommended this method for SV discovery in wheat (Fig. 4c,f and Supplementary Table 11).

A few earlier studies showed that high coverage of long-read sequencing was essential for the high-confidence discovery SVs [30, 62, 63]. For instance, ONT sequencing required at least 8X coverage of the human genome to achieve a precision rate of 0.8 [30]. HiFi sequencing, by contrast, has revolutionized the field with its highly accurate reads, significantly enhancing SV detection [50]. Nonetheless, the high cost of HiFi sequencing presents a challenge for its application at the population scale, particularly for the large and complex genomes such as wheat. Consequently, low-coverage HiFi sequencing becomes an appealing solution, enabling SV studies in large populations. Our examination of sequencing coverage revealed that, although some SV calls are missed, the precision of the SV call sets remains consistently high across coverage levels from 0.33X to 6.60X (Fig. 5). Considering that many genetic variants, including SVs, are shared across individuals, those SVs not detected in one individual could be captured from others within a large population. Thus, skim HiFi sequencing emerges as an ideal approach for investigating structural variations at the population level.

## Conclusion

Our study presented the first performance evaluation of leading SV aligner and caller using HiFi reads in plants. By introducing a widely applicable workflow for evaluating SV detection algorithms, we found that 1) a tailored approach that leverages the best aligner and caller for each SV type could improve the accuracy of SV discovery; 2) compared to the ensemble approach, the individual method with the highest F-score, can perform as well or better than the ensemble approach with much less computational resources; 3) low-coverage HiFi sequencing can provide high precision in SV calling, suggesting that skim HiFi sequencing is practical for large-scale, population-level SV studies. Taken together, the insights from this study are anticipated to facilitate studies of SVs’ functionality in wheat and many other plants.

## Methods

### Wheat accessions and PacBio HiFi sequencing

Three samples (AABBDD, AABB from *Triticum*; DD from *Aegilops*) were collected and planted in growth chambers. The tender leaves were divided into two equal parts, one half for next-generation sequencing (NGS) on the Illumina system and the other half for HiFi sequencing on the PacBio Sequel II system.

### Construction of the sequence-resolved candidate SV sets

#### Reads alignment

The HiFi reads were mapped to the corresponding wheat reference genome (IWGSC RefSeq v1.0) [40] using long-reads aligner (pbmm2 (version 1.5.0), NGMLR (version 0.2.7) [37], Winnowmap2 (version 2.0.3) [32], and Minimap2 (version 2.2.3) [64] with default parameters, respectively. The bam files were filtered and sorted using SAMtools (version 1.9) [65]. The detailed parameters were listed in Supplementary Table 10.

#### SV calling pipeline

SV calling was performed following the recommended parameters for HiFi reads using pbsv (version 2.3.0), cuteSV (version 1.0.9) [34], SVIM (version 1.4.2) [35], SVDSS (version 1.0.5) [36], and Sniffles2 (version 2.2) [37]). The minimal support read = 1 was set for SV calling due to the highly accurate HiFi reads. The detailed parameters were listed in Supplementary Table 10.

#### Candidate SV sets filter

SVs for 20 ACCs presenting the following conditions were retained: (1) SV length ≥ 50 bp; (2) minimal support long-read ≥ 1; (3) SVs passing the quality filters suggested by callers (flag PASS).

### The SV truth set construction

We deeply re-sequenced (14-25×) three wheat accessions to validate the results of each 20 ACCs through utilizing the discordant alignment and depth features. More details were provided below.

#### Read depth method for deletion truth set

For every given deletion from the above 20 candidate SV sets, read depth was first calculated by NGS data with mosdepth (version 0.2.6) [66]and the discordant alignment features were collected using SAMtools (FLAG 1294) [65] and the script “extractSplitReads_BwaMem” developed by lumpy-sv [67]. To obtain the more accurate truth set, we selected “Depth = 0” as the golden standard. Combining with the discordant alignment features, every given deletion was evaluated to determine if they were true. All true deletions, a higher resolution for breakpoints, obtained by the above method were merged and formed the deletion truth set.

#### Paragraph genotyping method for insertion truth set

We used Paragraph, an accurate genotyping tool for short-read sequencing data, to validate the insertion event that had been mined by 20 ACCs. Considering the record difference of position for the same insertion among SV callers, all true insertions obtained by Paragraph were merged using SURVIVOR [68] with 2 bp as the maximum allowed merging distance and formed the base-level insertion truth set.

Combined with the above results, the SV truth set was developed for further analysis.

### Validation of SV truth set by genome assembly, PCR amplification and Sanger sequencing

#### Genome assembly and assembly-based SV calling pipeline

We first performed *de novo* genome assembly for DD sample by HiCanu (version hicanu_rc) [46] using accurate long HiFi reads. SVMU (Structural Variants from MUMmer) was used to identify deletion and insertion via alignment of D-subgenome (from Chinese Spring reference) and DD contig sequence [69]. More precision results were obtained by Assemblytics [70] after MUMmer (version 4.0.0) [71]and LAST (version 1066) [72] alignment.

#### The comparison between the SV truth set and assembly-based SV dataset

The SV truth set was first filtered by the alignment regions obtained by minimap2, NGMLR and pbmm2 because the uncomplete genome (∼7X HiFi read data) could not cover the whole genome. Then, the resulting truth set was compared with the assembly-based SV dataset. For deletion, the deletion truth set was in comparison with MUMmer-SVMU pipeline using bedtools [73]. For insertion, the presence of at least two datasets from MUMmer-SVMU, MUMmer-Assemblytics and LAST-Assemblytics pipelines was in comparison with the insertion truth set.

#### PCR amplification and Sanger sequencing

PrimerServer was used to design genome-specific PCR primers for randomly selected SVs with default parameters [74]. The isolation of genome DNA for Chinese Spring and DD sample were performed based on the conventional cetyl trimethylammonium bromide (CTAB) method. PCR amplification products for each SV were validated by electrophoresis and used to perform Sanger sequencing using 3730*xl* DNA Analyzer.

### Evaluation of the ACCs and ensemble approach

The performance for 20 ACCs was assessed against the SV truth set using surpyvor (version 0.6.0) [30], a powerful tool for calculating precision-recall-F-score metrics. We further performed 20 times SRS (simple random sample) without replacement of size *n* (*n* = 2-20) from 20 ACCs and obtained a total of 1,600 ensemble SV sets through intersecting and uniting SV call sets of ACCs, while combining deletions and insertions, respectively. The intersection and union SV call sets were merged using SURVIVOR and evaluated against SV truth set with surpyvor. The precision, recall, and F-score were calculated as follows:

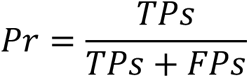

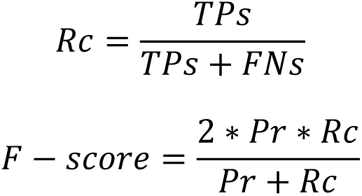

### The performance of low-coverage HiFi sequencing for SV calling

To study the relationship between the sequence coverage and performance, we randomly downsampled the sequencing data of the DD sample with 20 gradients from 5% (0.33×) to 100% (6.60×) using SAMtools. Each sample was performed and evaluated against the SV truth set after SV calling and filter with the above method.

## Availability of data and materials

The raw sequence data were deposited in the Genome Sequence Archive (GSA) (https://ngdc.cncb.ac.cn/gsa/) under accession numbers PRJCA005995.

## Authors’ contributions

FL designed and supervised the research. ZZ, JZ, CY and FL developed the general schematic workflow and performed data analysis. LK, SX, JX, YG and ZN helped with data analysis. XQ and BN performed PCR experiments. CY, AB, XZ, DX and JW collected plant materials. ZZ, JZ, and FL wrote the manuscript. All authors discussed the results and commented on the manuscript.

## Acknowledgments

We thank all members for their valuable comments in Fei Lu’s group, Institute of Genetics and Developmental Biology, Chinese Academy of Sciences.

## Reference

1. Alkan C, Coe BP, Eichler EE. Genome structural variation discovery and genotyping. Nat Rev Genet. 2011;12:363–76.

2. Rabanus-Wallace MT, Hackauf B, Mascher M, Lux T, Wicker T, Gundlach H, et al. Chromosome-scale genome assembly provides insights into rye biology, evolution and agronomic potential. Nature Genetics. 2021;53:564–73.

3. Pang AW, MacDonald JR, Pinto D, Wei J, Rafiq MA, Conrad DF, et al. Towards a comprehensive structural variation map of an individual human genome. Genome Biol. 2010;11:1–14.

4. Mahmoud M, Gobet N, Cruz-Dávalos DI, Mounier N, Dessimoz C, Sedlazeck FJ. Structural variant calling: The long and the short of it. Genome Biol. 2019;20:1–14.

5. Wendel JF, Jackson SA, Meyers BC, Wing RA. Evolution of plant genome architecture. Genome Biol. 2016;17:1–14.

6. Fu H, Dooner HK. Intraspecific violation of genetic colinearity and its implications in maize. Proceedings of the National Academy of Sciences. 2002;99:9573–8.

7. Brunner S, Fengler K, Morgante M, Tingey S, Rafalski A. Evolution of DNA Sequence Nonhomologies among Maize Inbreds. The Plant Cell Online. 2005;17:343–60.

8. Wang Q, Dooner HK. Remarkable variation in maize genome structure inferred from haplotype diversity at the bz locus. Proc Natl Acad Sci U S A. 2006;103:17644–9.

9. Alonge M, Wang X, Benoit M, Soyk S, Pereira L, Zhang L, et al. Major Impacts of Widespread Structural Variation on Gene Expression and Crop Improvement in Tomato. Cell. 2020;182:145–161.e23.

10. Zhang X, Zhu Y, Kremling KAG, Romay MC, Bukowski R, Sun Q, et al. Genome-wide analysis of deletions in maize population reveals abundant genetic diversity and functional impact. Theoretical and Applied Genetics. 2021;135:273–90.

11. Lu F, Romay MC, Glaubitz JC, Bradbury PJ, Elshire RJ, Wang T, et al. High-resolution genetic mapping of maize pan-genome sequence anchors. Nat Commun. 2015;6:6914.

12. Fuentes RR, Chebotarov D, Duitama J, Smith S, Hoz JFD la, Mohiyuddin M, et al. Structural variants in 3000 rice genomes. Genome Res. 2019;29:870–80.

13. Zhou Y, Minio A, Massonnet M, Solares E, Lv Y, Beridze T, et al. The population genetics of structural variants in grapevine domestication. Nat Plants. 2019;5:965–79.

14. Todesco M, Owens GL, Bercovich N, Légaré JS, Soudi S, Burge DO, et al. Massive haplotypes underlie ecotypic differentiation in sunflowers. Nature. 2020;584:602–7.

15. Castelletti S, Tuberosa R, Pindo M, Salvi S. A MITE transposon insertion is associated with differential methylation at the maize flowering time QTL Vgt1. G3 (Bethesda). 2014;4:805–12.

16. Yang Q, Li Z, Li W, Ku L, Wang C, Ye J, et al. CACTA-like transposable element in ZmCCT attenuated photoperiod sensitivity and accelerated the postdomestication spread of maize. Proceedings of the National Academy of Sciences. 2013;110:16969–74.

17. Huang C, Sun H, Xu D, Chen Q, Liang Y, Wang X, et al. ZmCCT9 enhances maize adaptation to higher latitudes. Proc Natl Acad Sci U S A. 2018;115:E334–41.

18. Liu J, Chen J, Zheng X, Wu F, Lin Q, Heng Y, et al. GW5 acts in the brassinosteroid signalling pathway to regulate grain width and weight in rice. Nature Plants 2017 3:5. 2017;3:1–7.

19. Wang Y, Xiong G, Hu J, Jiang L, Yu H, Xu J, et al. Copy number variation at the GL7 locus contributes to grain size diversity in rice. Nature Genetics 2015 47:8. 2015;47:944–8.

20. Nilsen KT, Walkowiak S, Xiang D, Gao P, Quilichini TD, Willick IR, et al. Copy number variation of TdDof controls solid-stemmed architecture in wheat. Proceedings of the National Academy of Sciences. 2020;117:28708–18.

21. Tikunov YM, Molthoff J, Vos RCH de, Beekwilder J, Houwelingen A van, Hooft JJJ van der, et al. NON-SMOKY GLYCOSYLTRANSFERASE1 Prevents the Release of Smoky Aroma from Tomato Fruit. Plant Cell. 2013;25:3067.

22. Liu Z, Roberts R, Mercer TR, Xu J, Sedlazeck FJ, Tong W. Towards accurate and reliable resolution of structural variants for clinical diagnosis. Genome Biol. 2022;23:1–25.

23. Chaisson MJP, Sanders AD, Zhao X, Malhotra A, Porubsky D, Rausch T, et al. Multi-platform discovery of haplotype-resolved structural variation in human genomes. Nat Commun. 2019;10:1–16.

24. Yuan Y, Bayer PE, Batley J, Edwards D. Current status of structural variation studies in plants. Plant Biotechnol J. 2021;19:2153–63.

25. Shi J, Tian Z, Lai J, Huang X. Plant pan-genomics and its applications. Mol Plant. 2023;16:168–86.

26. De Coster W, Van Broeckhoven C. Newest Methods for Detecting Structural Variations. Trends Biotechnol. 2019;37:973–82.

27. Logsdon GA, Vollger MR, Eichler EE. Long-read human genome sequencing and its applications. Nat Rev Genet. 2020;21:597–614.

28. Ho SS, Urban AE, Mills RE. Structural variation in the sequencing era. Nat Rev Genet. 2020;21:171–89.

29. Zhou A, Lin T, Xing J. Evaluating nanopore sequencing data processing pipelines for structural variation identification. Genome Biol. 2019;20:1–13.

30. Coster W De, Rijk P De, Roeck A De, Pooter T De, D’Hert S, Strazisar M, et al. Structural variants identified by Oxford Nanopore PromethION sequencing of the human genome. Genome Res. 2019;29:gr.244939.118.

31. Sedlazeck FJ, Rescheneder P, Smolka M, Fang H, Nattestad M, Haeseler A von, et al. Accurate detection of complex structural variations using single molecule sequencing. Nat Methods. 2018;15:461.

32. Jain C, Rhie A, Hansen NF, Koren S, Phillippy AM. Long-read mapping to repetitive reference sequences using Winnowmap2. Nat Methods. 2022;19:705–10.

33. Li H. New strategies to improve minimap2 alignment accuracy. Bioinformatics. 2021; October:1–3.

34. Cheng H, Concepcion GT, Feng X, Zhang H, Li H. Haplotype-resolved de novo assembly with phased assembly graphs. 2020. 10.1038/s41592-020-01056-5.

35. D Heller MV. SVIM: structural variant identification using mapped long reads. Bioinformatics. 2019;35:2907–15.

36. Denti L, Khorsand P, Bonizzoni P, Hormozdiari F, Chikhi R. SVDSS: structural variation discovery in hard-to-call genomic regions using sample-specific strings from accurate long reads. Nat Methods. 2022;20:550–8.

37. Sedlazeck FJ, Rescheneder P, Smolka M, Fang H, Nattestad M, Haeseler A von, et al. Accurate detection of complex structural variations using single molecule sequencing. Nat Methods. 2018;15:461.

38. Sahlin K, Baudeau T, Cazaux B, Marchet C. A survey of mapping algorithms in the long-reads era. Genome Biol. 2023;24:1–23.

39. Ahsan MU, Liu Q, Perdomo JE, Fang L, Wang K. A survey of algorithms for the detection of genomic structural variants from long-read sequencing data. Nat Methods. 2023;20:1143–58.

40. Appels R, Eversole K, Feuillet C, Keller B, Rogers J, Stein N, et al. Shifting the limits in wheat research and breeding using a fully annotated reference genome. Science. 2018;361:eaar7191.

41. Zook JM, Hansen NF, Olson ND, Chapman L, Mullikin JC, Xiao C, et al. A robust benchmark for detection of germline large deletions and insertions. Nat Biotechnol. 2020;38:1347–55.

42. Collins RL, Brand H, Karczewski KJ, Zhao X, Alföldi J, Francioli LC, et al. A structural variation reference for medical and population genetics. Nature. 2020;581:444–51.

43. Zhao X, Guo Y, Kang L, Bi A, Xu D, Zhang Z, et al. Population genomics unravels the Holocene history of Triticum-Aegilops species. bioRxiv. 2022;:2022.04.07.487499.

44. Zhou Y, Zhao X, Li Y, Xu J, Bi A, Kang L, et al. Triticum population sequencing provides insights into wheat adaptation. Nat Genet. 2020;52:1412–22.

45. S Chen PKEDRSRPFSMKDBMSFSME. Paragraph: A graph-based structural variant genotyper for short-read sequence data. bioRxiv. 2019;24:635011.

46. Nurk S, Walenz BP, Rhie A, Vollger MR, Logsdon GA, Grothe R, et al. HiCanu: accurate assembly of segmental duplications, satellites, and allelic variants from high-fidelity long reads. Genome Res. 2020;30:gr.263566.120.

47. Cameron DL, Di Stefano L, Papenfuss AT. Comprehensive evaluation and characterisation of short read general-purpose structural variant calling software. Nat Commun. 2019;10:1–11.

48. Kosugi S, Momozawa Y, Liu X, Terao C, Kubo M, Kamatani Y. Comprehensive evaluation of structural variation detection algorithms for whole genome sequencing. Genome Biol. 2019;20:8–11.

49. Nielsen R, Paul JS, Albrechtsen A, Song YS. Genotype and SNP calling from next-generation sequencing data. Nat Rev Genet. 2011;12:443–51.

50. Wenger AM, Peluso P, Rowell WJ, Chang PC, Hall RJ, Concepcion GT, et al. Accurate circular consensus long-read sequencing improves variant detection and assembly of a human genome. Nat Biotechnol. 2019;37:1155–62.

51. Liu Y, Du H, Li P, Shen Y, Peng H, Liu S, et al. Pan-Genome of Wild and Cultivated Soybeans. Cell. 2020;182:162–176.e13.

52. Hufford MB, Seetharam AS, Woodhouse MR, Chougule KM, Ou S, Liu J, et al. De novo assembly, annotation, and comparative analysis of 26 diverse maize genomes. Science. 2021;373:655–62.

53. Qin P, Lu H, Du H, Wang H, Chen W, Chen Z, et al. Pan-genome analysis of 33 genetically diverse rice accessions reveals hidden genomic variations. Cell. 2021;:1–17.

54. Zhou Y, Zhang Z, Bao Z, Li H, Lyu Y, Zan Y, et al. Graph pangenome captures missing heritability and empowers tomato breeding. Nature. 2022;606:527–34.

55. Tang D, Jia Y, Zhang J, Li H, Cheng L, Wang P, et al. Genome evolution and diversity of wild and cultivated potatoes. Nature. 2022;606:535–41.

56. Li N, He Q, Wang J, Wang B, Zhao J, Huang S, et al. Super-pangenome analyses highlight genomic diversity and structural variation across wild and cultivated tomato species. Nat Genet. 2023;55:852–60.

57. Huang Y, He J, Xu Y, Zheng W, Wang S, Chen P, et al. Pangenome analysis provides insight into the evolution of the orange subfamily and a key gene for citric acid accumulation in citrus fruits. Nat Genet. 2023;55:1964–75.

58. He Q, Tang S, Zhi H, Chen J, Zhang J, Liang H, et al. A graph-based genome and pangenome variation of the model plant Setaria. Nat Genet. 2023;55:1232–42.

59. Chen J, Liu Y, Liu M, Guo W, Wang Y, He Q, et al. Pangenome analysis reveals genomic variations associated with domestication traits in broomcorn millet. Nat Genet. 2023. 10.1038/s41588-023-01571-z.

60. De Coster W, Weissensteiner MH, Sedlazeck FJ. Towards population-scale long-read sequencing. Nat Rev Genet. 2021;22:572–87.

61. Olson ND, Wagner J, Dwarshuis N, Miga KH, Sedlazeck FJ, Salit M, et al. Variant calling and benchmarking in an era of complete human genome sequences. Nat Rev Genet. 2023;24:464–83.

62. Sedlazeck FJ, Rescheneder P, Smolka M, Fang H, Nattestad M, Von Haeseler A, et al. Accurate detection of complex structural variations using single-molecule sequencing. Nat Methods. 2018;15:461–8.

63. Tham CY, Tirado-Magallanes R, Goh Y, Fullwood MJ, Koh BTH, Wang W, et al. NanoVar: Accurate characterization of patients’ genomic structural variants using low-depth nanopore sequencing. Genome Biol. 2020;21:1–15.

64. Li H. New strategies to improve minimap2 alignment accuracy. Bioinformatics. 2021; October:1–3.

65. Li H, Handsaker B, Wysoker A, Fennell T, Ruan J, Homer N, et al. The Sequence Alignment/Map format and SAMtools. Bioinformatics. 2009;25:2078–9.

66. Pedersen BS, Quinlan AR. Mosdepth: Quick coverage calculation for genomes and exomes. Bioinformatics. 2018;34:867–8.

67. Layer RM, Chiang C, Quinlan AR, Hall IM. LUMPY: A probabilistic framework for structural variant discovery. Genome Biol. 2014;15:1–19.

68. Jeffares DC, Jolly C, Hoti M, Speed D, Shaw L, Rallis C, et al. Transient structural variations have strong effects on quantitative traits and reproductive isolation in fission yeast. Nature Communications 2017 8:1. 2017;8:1–11.

69. Chakraborty M, Emerson JJ, Macdonald SJ, Long AD. Structural variants exhibit widespread allelic heterogeneity and shape variation in complex traits. Nat Commun. 2019;10.

70. Nattestad M, Schatz MC. Assemblytics: A web analytics tool for the detection of variants from an assembly. Bioinformatics. 2016;32:3021–3.

71. Marçais G, Delcher AL, Phillippy AM, Coston R, Salzberg SL, Zimin A. MUMmer4: A fast and versatile genome alignment system. PLoS Comput Biol. 2018;14:e1005944.

72. Kiełbasa SM, Wan R, Sato K, Horton P, Frith MC. Adaptive seeds tame genomic sequence comparison. Genome Res. 2011;21:487–93.

73. Quinlan AR, Hall IM. BEDTools: A flexible suite of utilities for comparing genomic features. Bioinformatics. 2010;26:841–2.

74. Zhu T, Liang C, Meng Z, Li Y, Wu Y, Guo S, et al. PrimerServer: a high-throughput primer design and specificity-checking platform. bioRxiv. 2017. 10.1101/181941.

