## Supplementary figures for "Structural variation discovery in wheat using PacBio high-fidelity sequencing": Supplementary_figures.docx

**Title:**

**File contents:**

Supplementary Figures 1-10

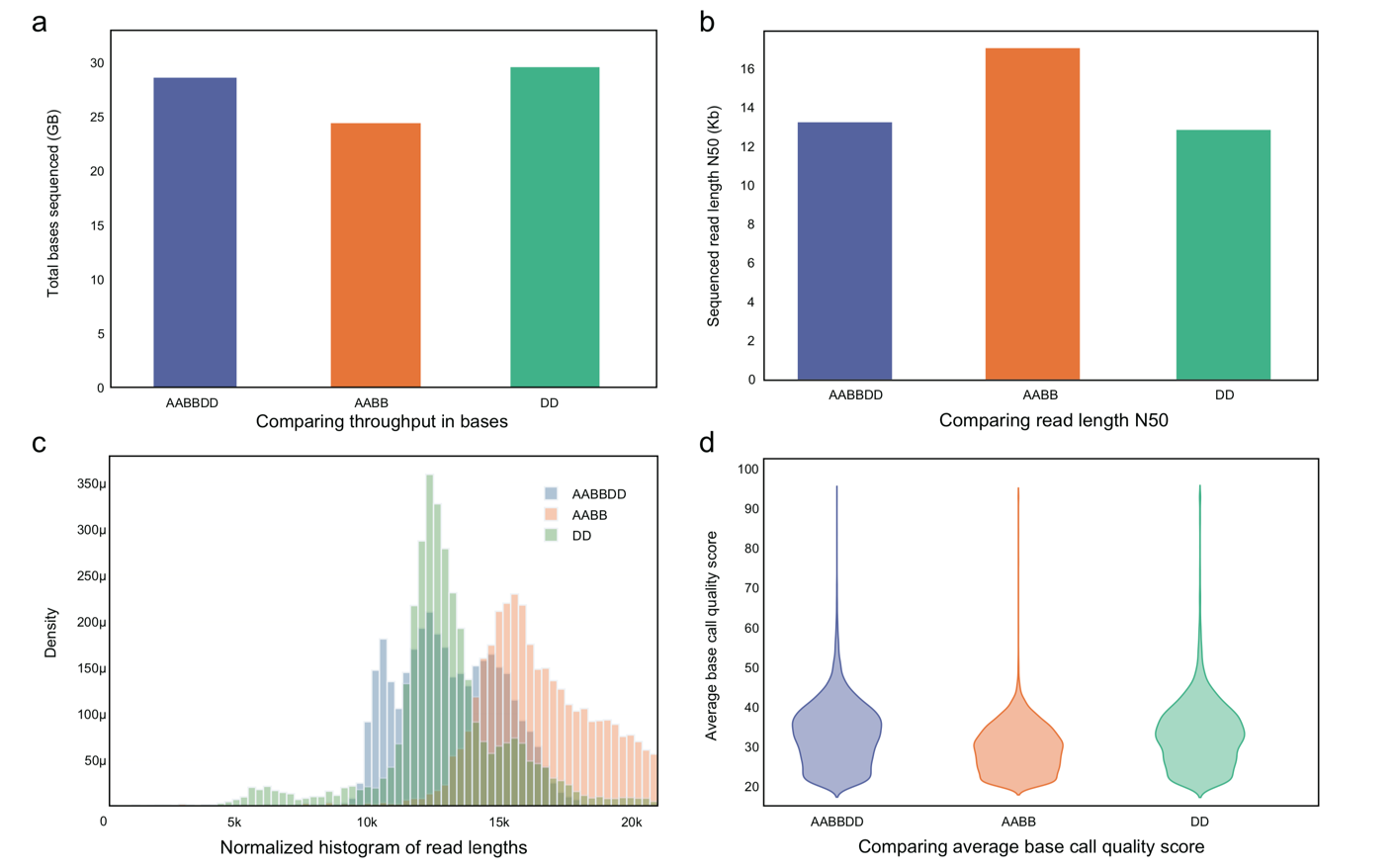

**Supplementary Figure 1.** Summary of AABBDD, AABB, DD in this study. **a - d** showed total output filed (Gb), the length of HiFi reads N50, the distribution of HiFi reads length, and the average base quality score.

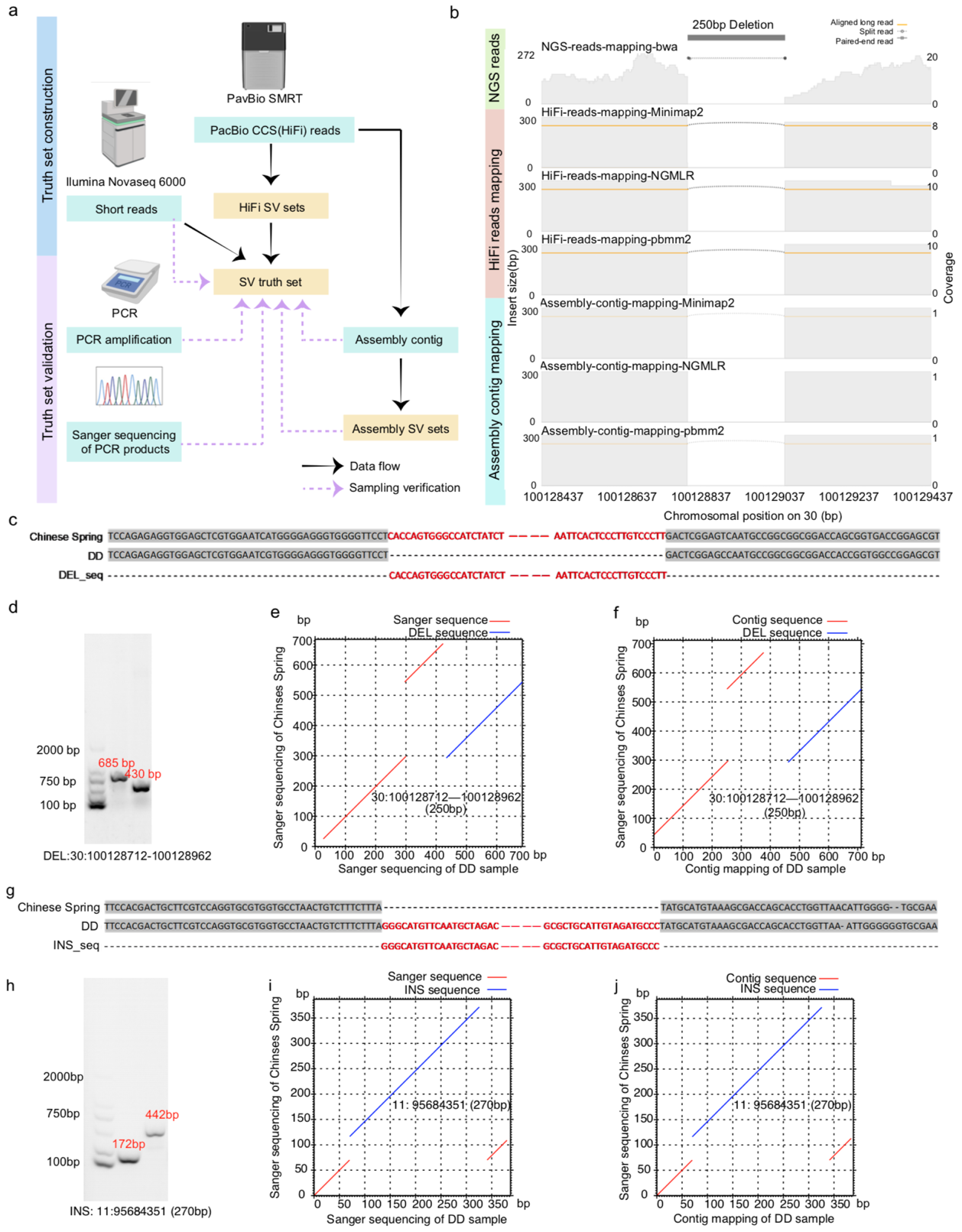

**Supplementary Figure 2.** Validation of the SV truth set. An example validated a 250-bp deletion event by integrating a total of 10 methods based on PCR amplification, sanger sequencing, NGS reads mapping, HiFi reads mapping, contig mapping and whole genome alignment (a-f). Another example of 270 bp insertion supported by PCR amplification, sanger sequencing and genome assembly validated the high-confidence insertion true set (g-j).

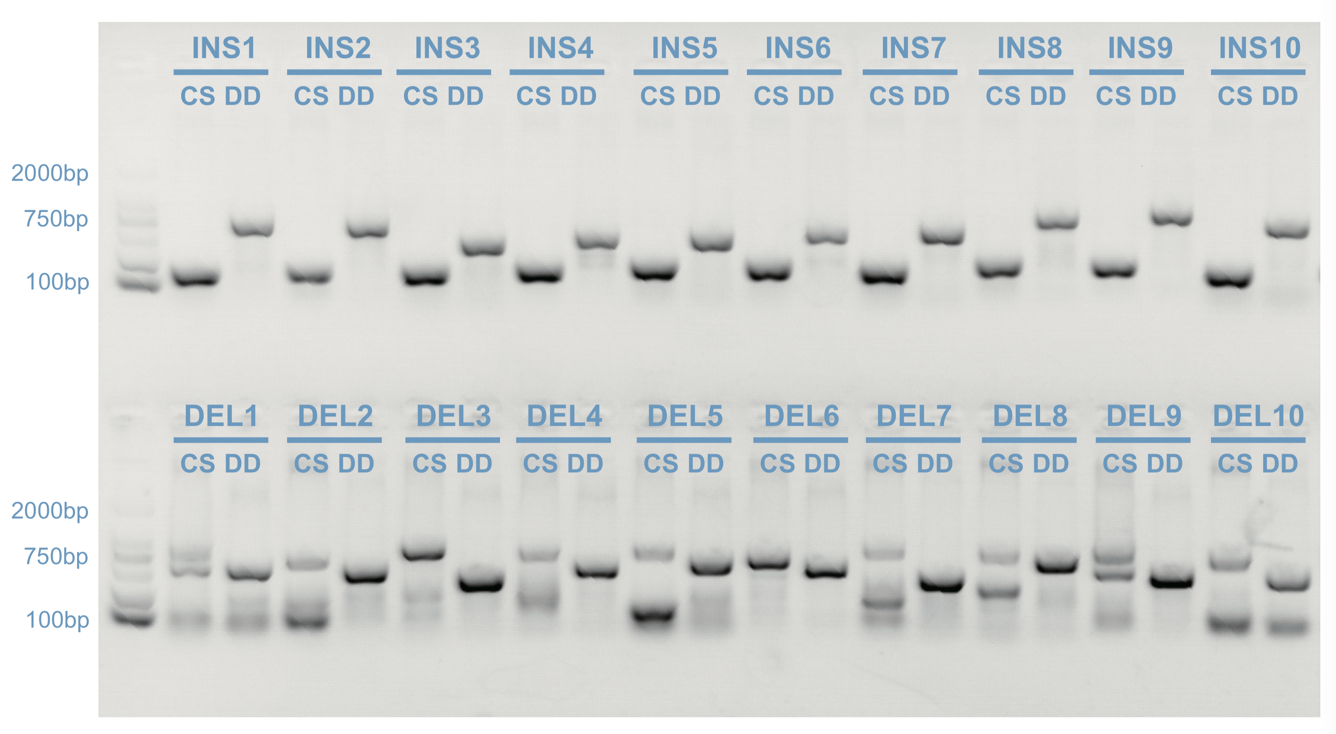

**Supplementary** **Figure 3.** PCR amplification validated the high-confidence SV true set. A total of 20 randomly selected SVs from the base-level true set were validated by PCR, reflecting the reliability of the SV true set. Interestingly, DEL1 and DEL9 reported two clear bands, present/absent of the locus, representing a heterozygous locus.

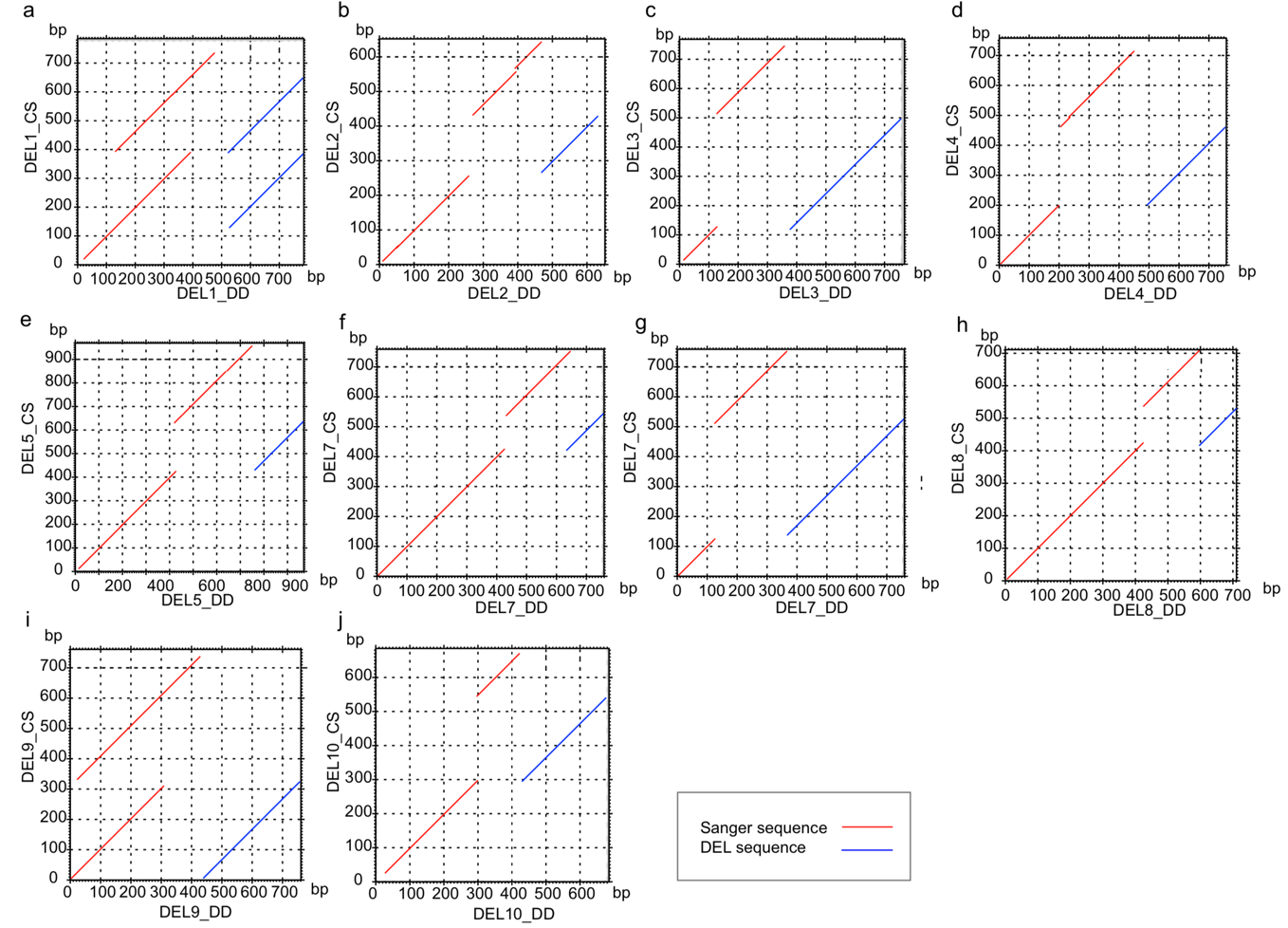

**Supplementary** **Figure 4.** Sequence collinearity for deletion between Chinese Spring and DD sample via sanger sequencing.

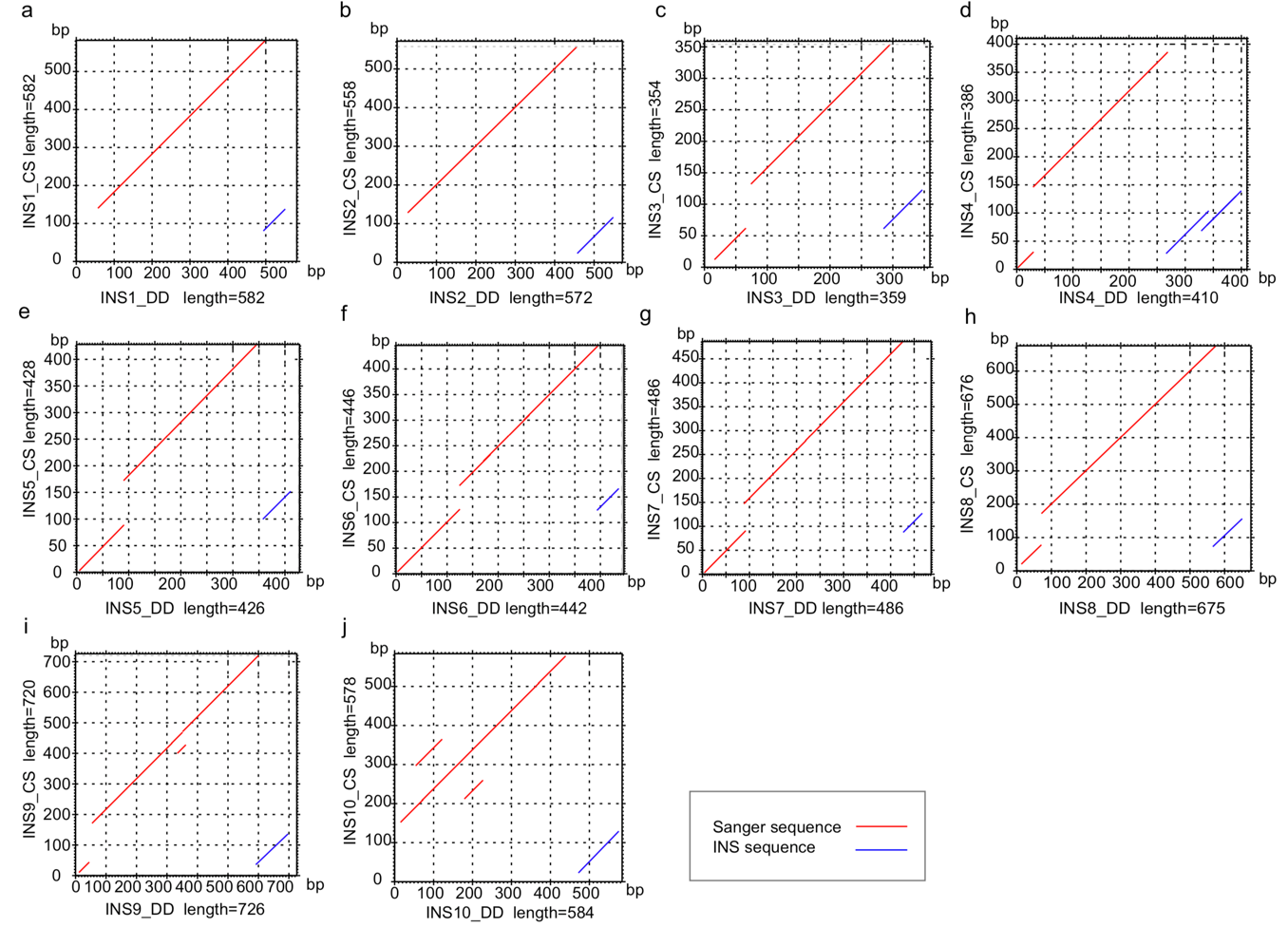

**Supplementary Figure 5.** Sequence collinearity for insertion between Chinese Spring and DD sample via sanger sequencing.

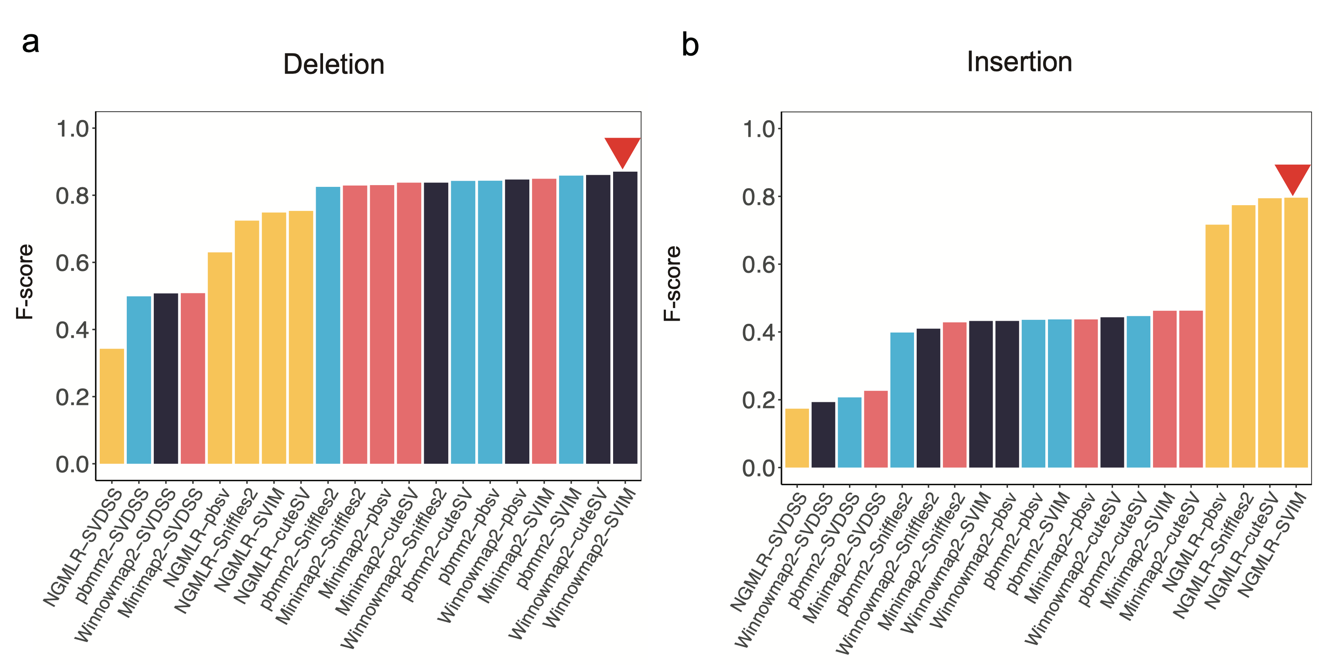

**Supplementary Figure 6.** F-score comparison of 20 ACCs against the SV truth set. The inverse red triangle denoted the maximum F-measure **(a, b)**. Aligner Minimap2, NGMLR, Winnowmap2, and pbmm2 were tagged with red, orange, black, and blue, respectively.

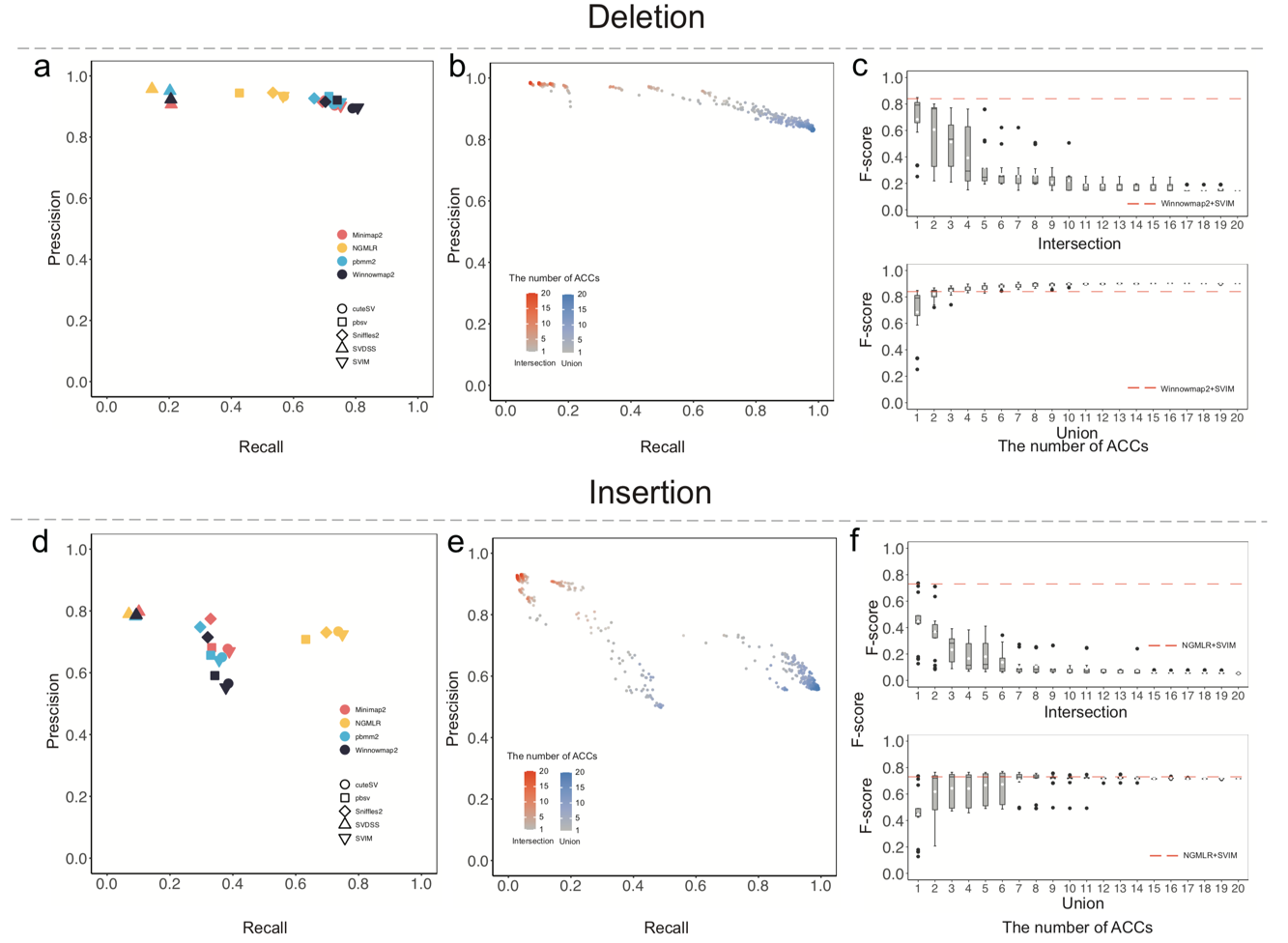

**Supplementary Figure 7.** Comprehensive evaluation of individual ACCs and the ensemble approach in DD sample. The precision-recall graph of 20 ACCs against the SV truth set (a, d). The performance of 800 ensemble SV sets through intersecting and uniting SV call sets of ACCs, while combining deletions and insertions, respectively (b, e). The F-score comparison between 800 ensemble SV sets and the highest F-score in 20 ACCs (c, f). The highest F-score of 20 ACCs was represented by red dashed line.

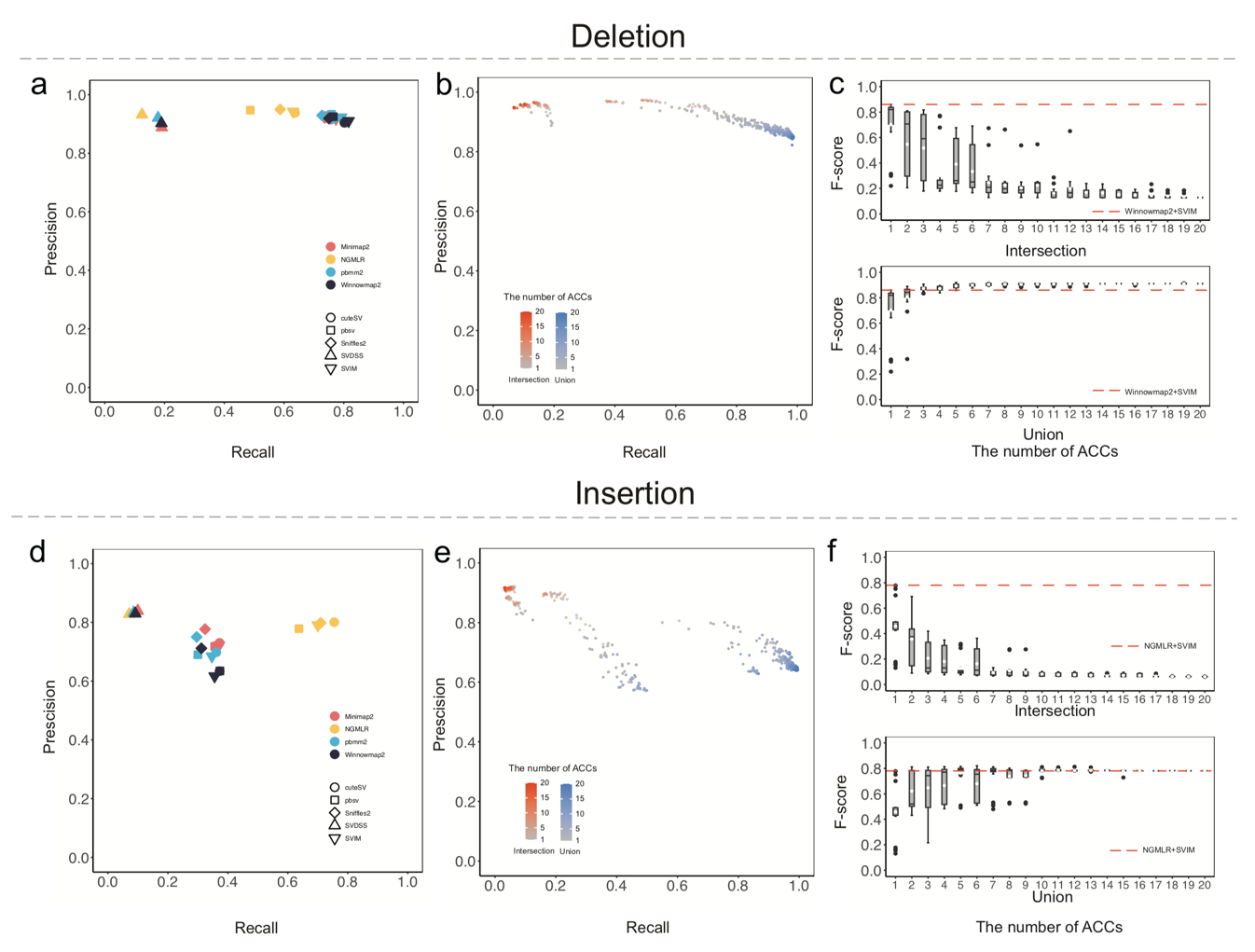

**Supplementary** **Figure 8.** Comprehensive evaluation of individual ACCs and the ensemble approach in AABB sample. The precision-recall graph of 20 ACCs against the SV truth set (a, d). The performance of 800 ensemble SV sets through intersecting and uniting SV call sets of ACCs, while combining deletions and insertions, respectively (b, e). The F-score comparison between 800 ensemble SV sets and the highest F-score in 20 ACCs (c, f). The highest F-score of 20 ACCs was represented by red dashed line.

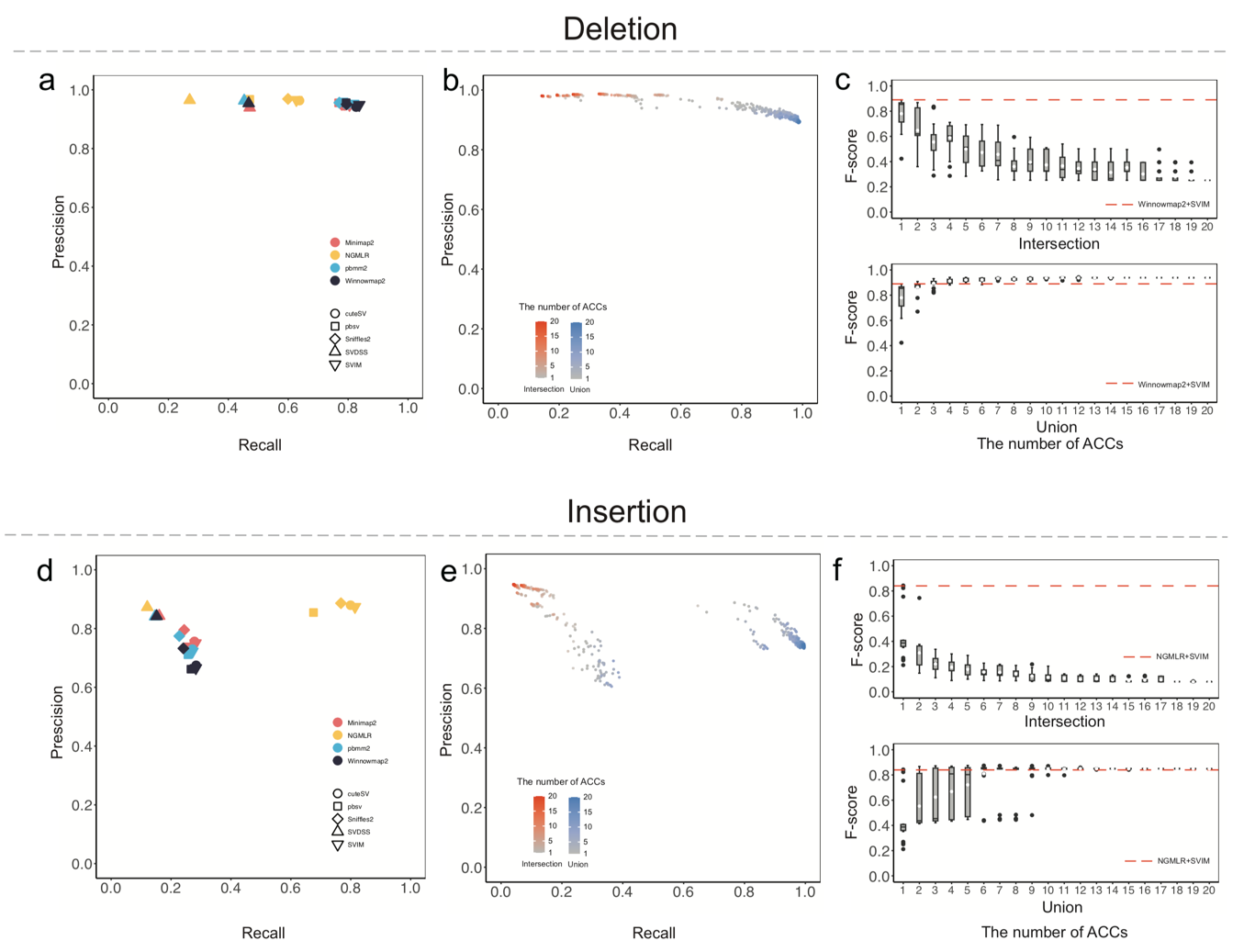

**Supplementary** **Figure 9.**  Comprehensive evaluation of individual ACCs and the ensemble approach in AABBDD sample. The precision-recall graph of 20 ACCs against the SV truth set (a, d). The performance of 800 ensemble SV sets through intersecting and uniting SV call sets of ACCs, while combining deletions and insertions, respectively (b, e). The F-score comparison between 800 ensemble SV sets and the highest F-score in 20 ACCs (c, f). The highest F-score of 20 ACCs was represented by red dashed line.

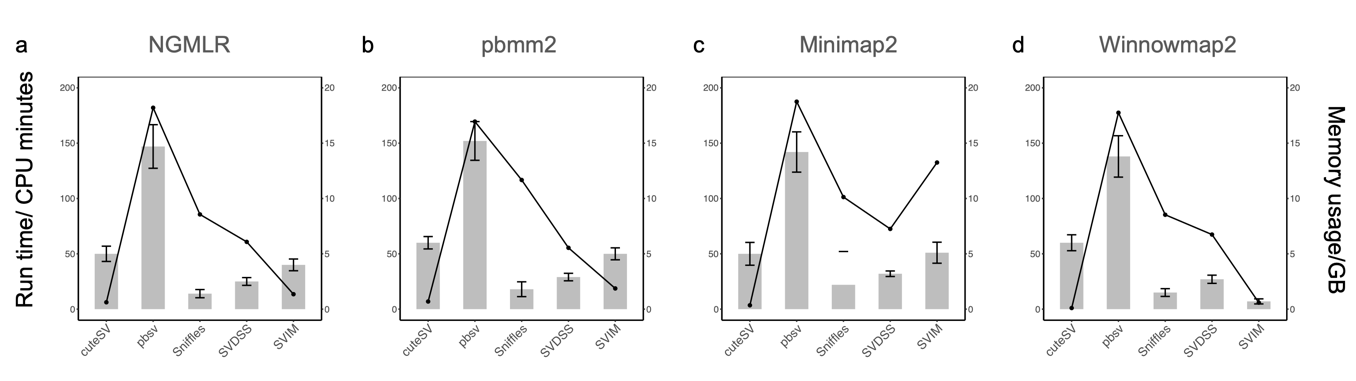

**Supplementary** **Figure 10.** Run time and memory consumption for SV detection algorithms. The occupancy of computing resources, an important factor considered by users including run time and average memory usage, was then examined using 20 threads in the AABBDD genome. For run time, aligner pbmm2, Minimap2, and Winnowmap2 generally processed the same datasets (~30Gb) with 5-7 times less runtime relative to NGMLR (951.0 & 616.8 & 2059.6 vs. 6727.5 CPU minutes), and callers pbsv took a long time for SV detection than the other three callers. For average memory usage, aligners occupied relatively high memory (32-45 G). In addition, callers cuteSV, Sniffles, SVDSS, and SVIM used a similar memory (≤ 6 G), and pbsv required a little more memory (~15 G) **(a-c)**. Overall, the impact on computing resources was mainly concentrated in the aligner, not the SV caller.
